# Bioinformatic analysis of metastasis-associated metabolic landscape reveals an oncogenic role for the transsulfuration pathway

**DOI:** 10.64898/2025.12.07.692828

**Authors:** Jonathan K. Yan, Ying Yang, Wenqi Wang

## Abstract

Cancer metastasis is a leading cause of cancer-related deaths, while its underlying mechanisms remain incompletely understood. To colonize distant organs, cancer cells reprogram their metabolism to adapt to diverse environmental challenges. Therefore, elucidating the metabolic pathways that drive cancer metastasis will uncover novel biomarkers and therapeutic targets. In this study, we integrated published datasets and systematically analyzed metabolites across multiple cancer cell lines. This large-scale bioinformatic analysis revealed distinct metabolites and metabolic pathways associated with organ-specific metastasis, and underscored the crucial role of tissue of origin in shaping the metabolic landscape of metastatic tumors. Notably, the transsulfuration pathway (also known as the cysteine and methionine metabolism) was strongly enriched in cancer cells with high metastatic potential. We validated this finding in pancreatic cancer, where the pathway enzyme cystathionine β-synthase (CBS) and its metabolic products were highly expressed in metastatic cancer cells. Targeting the transsulfuration pathway either by methionine deprivation or pharmacological inhibition of CBS significantly impaired the migration and invasion of metastatic pancreatic cancer cells. Taken together, our study not only provides a global view of the altered metabolic landscape in metastasis, but also identifies the transsulfuration pathway as an oncogenic driver and a therapeutic target for pancreatic cancer metastasis.

## Introduction

Cancer is a leading cause of death worldwide [1], with cancer metastasis accounting for ∼90% of cancer-related mortalities [2]. While advances in treating primary tumors have improved patient outcome, effective therapeutic options for cancer metastasis remain limited.

Metastasis is the process by which a subset of cancer cells disseminates from the primary tumor and spreads to distant organs [3–5]. This complex process includes local tumor cell invasion, entry into the vasculature, exit from the circulation, and successful colonization at distant tissues. Metastatic cells must overcome a series of challenges, including acquiring motility, invading surrounding tissue, and surviving in circulation and distant organs [6–8].

Although only a small fraction of cancer cells can eventually succeed in establishing secondary tumors, metastasis is not a random event but rather a highly regulated process by both intrinsic tumor molecular features and the microenvironment of target organs. This concept was first proposed by Stephen Paget in 1889 through the “seed and soil” hypothesis, suggesting that metastatic cancer cells (the “seed”) selectively colonize organs with favorable microenvironments (the “soil”) [9]. Recent studies have shown that genetic mutations and epigenetic alterations play critical roles in shaping the metastatic potential of cancer cells [10–12]. Moreover, cancer cells undergo significant metabolic reprogramming to support their rapid growth and survival, driven by the genetic and epigenetic changes. For example, they often increase the uptake of nutrients, such as glucose and glutamine, to fuel proliferation [13, 14]. Additionally, cancer cells develop strategies to adapt to metabolic stresses, such as nutrients and oxygen deprivation in the tumor microenvironment [15]. While research has been focused on how metabolism regulates cancer proliferation and survival, only recently has attention turned to the role of metabolic reprogramming that enables metastatic cells to thrive in a foreign microenvironment (i.e., new “soil”) [16–19]. However, a systematic comparison of metabolic alterations between metastatic and non-metastatic cancer cells remains lacking.

In this study, we developed a strategy to systemically examine the association of cancer metabolism and metastasis by integrating two complementary datasets derived from the Cancer Cell Line Encyclopedia (CCLE) – the Metastasis Map (MetMap) [20] and the large-scale metabolomic profiling [21]. We found that metastasis-associated metabolic landscape exhibits organ-specific features, supporting the idea that diverse microenvironments require distinct metabolic adaptations. In addition, tumor tissue of origin influences the metabolic landscape of metastasis. Interestingly, we identified the transsulfuration pathway as a critical metabolic driver of metastasis. In pancreatic cancer, targeting the transsulfuration pathway, either through methionine deprivation or pharmacological inhibition, significantly suppressed the migration and invasion of metastatic cancer cells. Collectively, our study offers a comprehensive profile of the altered metabolic landscape in metastasis, revealing candidate biomarkers and targets for combatting metastatic cancers.

## Methods

### Bioinformatic analysis

Two datasets were used in this study. The first dataset examined the metastatic potential of 500 human cancer cell lines was determined using *in vivo* mouse tumor models combined with a DNA-based barcoding strategy [20]. This dataset comprises six metastatic tissue destinations – overall, bone, brain, lung, liver, and kidney, classified as follows: potential value ≤ −4 (non-metastatic), between −4 and −2 (weakly metastatic), and ≥ −2 (metastatic with higher confidence) [20]. For our analysis, cell lines with metastatic potential values ≤ −2 were classified as non-metastatic, and those with values > −2 were classified as metastatic. The second dataset profiled 225 metabolites across 928 cell lines from over 20 cancer types in the Cancer Cell Line Encyclopedia (CCLE) using liquid chromatography-mass spectrometry (LC-MS) [21]. These two datasets were integrated to identify cancer metastasis-associated metabolites and metabolic pathways.

Using R, metastatic potential values ≤ −2 were assigned “0” (non-metastatic) and values > −2 were assigned “1” (metastatic). Metabolite values from the second dataset were obtained through anti-logged process. The two datasets were then merged via inner join, connecting metabolite measurements with metastatic potential for cell lines present in both datasets, with a total of 479 cell lines **(Table S1)**. Similarly, datasets for organ-specific metastasis (bone, brain, liver, lung, and kidney) were integrated with metabolic profiles, resulting profile from 479 cell lines.

Data processing was performed using the MetaboAnalystR package [22]. *T*-test was used to assess correlations between individual metabolites and metastatic potential. Metabolites with a raw *p-*value of < 0.05 were considered significantly associated with metastasis. Volcano plots were generated using thresholds of *p* < 0.05 and fold change ≥ 1.2. Significant metabolites were then subjected to enrichment and pathway analyses to identify metabolic pathways associated with metastasis. To analyze metabolites across different metastasis sites (bone, brain, liver, lung, and kidney), UpSet plots were generated in R using the UpSetR package [23]. Enrichment analysis was performed for metabolites shared by three or more of the metastatic sites. Cell lines from five highly metastatic cancer types (breast, skin, ovarian, colon, and pancreatic cancer), were extracted, and volcano plots were generated for each tissue of origin **(Table S1)**. UpSet plots were also generated to illustrate significant metabolites shared across different metastatic sites from each tissue of origin.

### Cell culture and chemicals

PATU8988, SUIT2, HUPT3, PANC-1 and CAPAN2 were kindly provided by Dr. Christopher Halbrook (University of California, Irvine) and cultured in Dulbecco’s modified Eagle’s medium (DMEM, Corning, New York, NY, USA) containing 25 mM glucose and 4 mM L-glutamine. DMEM was supplemented with 10% fetal bovine serum (FBS, Gemini Bio-Products, Sacramento, CA, USA), 100 units/mL penicillin, and 100 μg/mL streptomycin (Sigma). All cells were cultured at 37°C with 5% CO_2_. Methionine-free medium was made with DMEM devoid of methionine and L-Cystine (Gibco, 21013-024) supplemented with 10% dialyzed FBS (dFBS) (Omega Scientific, Tarzana, CA, USA) and 2 mM L-glutamine (Omega Scientific, Tarzana, CA, USA). Complete DMEM was made by supplementing with 2 mM L-Glutamine and 10% dFBS.

Aminooxyacetic Acid (AOAA) and Epigallocatechin Gallate (EGCG) were purchased from Cayman Chemical (Ann Arbor, MI, USA).

### Metabolites measurement by gas chromatography–mass spectrometry (GC-MS)

PATU8988, SUIT2, HUPT3 and CAPAN2 cells were cultured in complete or methionine-free medium for 24 hours and washed with PBS and 80% methanol/water (HPLC grade) with norvaline added as an internal standard. The plate was transferred to a −80 _°_C freezer for 15 min to further inactivate enzymes. Cells were then harvested by a silicone scraper, and the whole-cell extract was transferred to a tube and centrifuged at 17000 g for 10 min at 4 _°_C. The supernatant was transferred into tubes and dried by speed vacuum. 50 µL of MOX (10 mg/ml in pyridine, 226904 Sigma) was added, and the mixture was incubated at 42 _°_C for 1 hour. After the samples were cooled down, 100 μl TBDMS (394882, Sigma) was added, and the samples were incubated at 70 _°_C for 1 hour. Samples were transferred to GC vials and analyzed by Agilent 7820 A chromatography and Agilent 5977B mass spectrometer. Results were processed by GraphPad Prism (v 9) software, where *t*-tests were performed to determine the statistical significance of the differences between the means of the analyzed groups.

### Wound healing assay

PATU8988 (8 × 10^5^ cells per well) and SUIT2 (6 × 10^5^ cells per well), PANC-1 (5 × 10^5^ cells per well) cells were seeded in 6-well plates. After 24 hours, the cells were scratched with a 200µL sterile pipette tip to create a cross-shaped pattern. Cells were washed with phosphate-buffered saline (PBS) (1×) to remove any detached cells. For methionine deprivation experiments, either complete DMEM with dFBS or methionine-free medium was added to the well. For the drug treatment, AOAA or EGCG was added to the DMEM to make the final concentration of 10 mM or 0.1 mM, respectively. Three wells for each condition were made. The plates were then placed under a microscope (EVOS Cell Imaging Systems) using the center cross as the reference mark. Four pictures of wounds were taken at the spots that are directly left, right, up, and down from the cross using 4× objective. The wound area was then calculated using the wound healing size plugin for Image J as described in the paper by Suarrez-Arnedo A. et.al. [24]. For propidium iodide (PI) staining, the stock solution (P4864, Sigma-Aldrich) was diluted to 1/100 solution/medium. Cells were incubated for five minutes and analyzed under microscope (EVOS Cell Imaging Systems).

### Cell invasion assay

Matrigel inserts (Corning, Bedford, MA, USA) were rehydrated according to the manufacturer’s instructions with control inserts placed into empty wells of a 6-well plate. SUIT2 cell suspensions were prepared with serum-free DMEM with or without methionine at 1.5 × 10^5^ cells/ml. DMEM containing 10% of FBS with or without methionine was added to the wells, and inserts were carefully transferred to avoid air bubble entrapment. Cells were seeded into the inserts (2 mL of 1.5 × 10^5^ cells for 6-well chambers) and incubated for 22 hours. The inserts were washed with PBS, and non-invading cells were removed by scrubbing the upper membrane surface with a cotton-tipped swab moistened with PBS. The invaded cells on the lower membrane surface were then fixed using 100% methanol for 20 minutes at room temperature. The membranes were washed with PBS, stained with 0.2% crystal violet stain solution for 10 minutes, and washed three times with distilled water.

### Western blotting

Cells were lysed on ice in lysis buffer (50 mM Tris-HCL [pH 7.4], 5 mM Sodium Fluoride, 5 mM Sodium Pyrophosphate, 1 mM EDTA, 1 mM EGTA, 250 mM Mannitol, 1% [v/v] Triton X-100) containing protease inhibitor complex (04693159001, Roche) and phosphatase inhibitor (1862495, Thermo fisher scientific). The protein concentrations were measured by using the BCA Assay kit (23225, Thermo Fisher Scientific). The immunoblotting was performed with the following antibodies: anti-CBS (14782, Cell Signaling, 1:1000 dilution), anti-CTH/CGL (GTX 113409, GeneTex, 1:1000 dilution), and anti-β-actin (A5441, Sigma-Aldrich, 1:5000 dilution).

### Statistics

Results are shown as means, and error bars represent standard deviation (s.d.). The unpaired two-tailed Student’s *t*-test was used to determine the statistical significance of differences (\**p* < 0.05, \*\**p* < 0.01, \*\*\**p* < 0.001, \*\*\*\**p* < 0.0001) using Microsoft Excel or GraphPad Prism (v 9) software.

## Results

### Bioinformatic analysis of cancer metastasis-related metabolic pathways

To identify metabolic pathways associated with metastasis, we integrated the published MetMap dataset [20] with metabolomics data from the Cancer Cell Line Encyclopedia (CCLE) [21], connecting the metabolic profiles of cancer cells to their metastatic potential. Metabolite levels were further compared between metastatic and non-metastatic cells (**Figure 1A**). Interestingly, metabolites, such as homocysteine, asparagine, phosphocreatine, and phosphogluconate were significantly enriched in metastatic cancer cells, whereas metabolites, including thymine, serine, and certain triglycerides (TAGs) were largely reduced (**Figure 1A** and Table S2).

**Figure 1.**
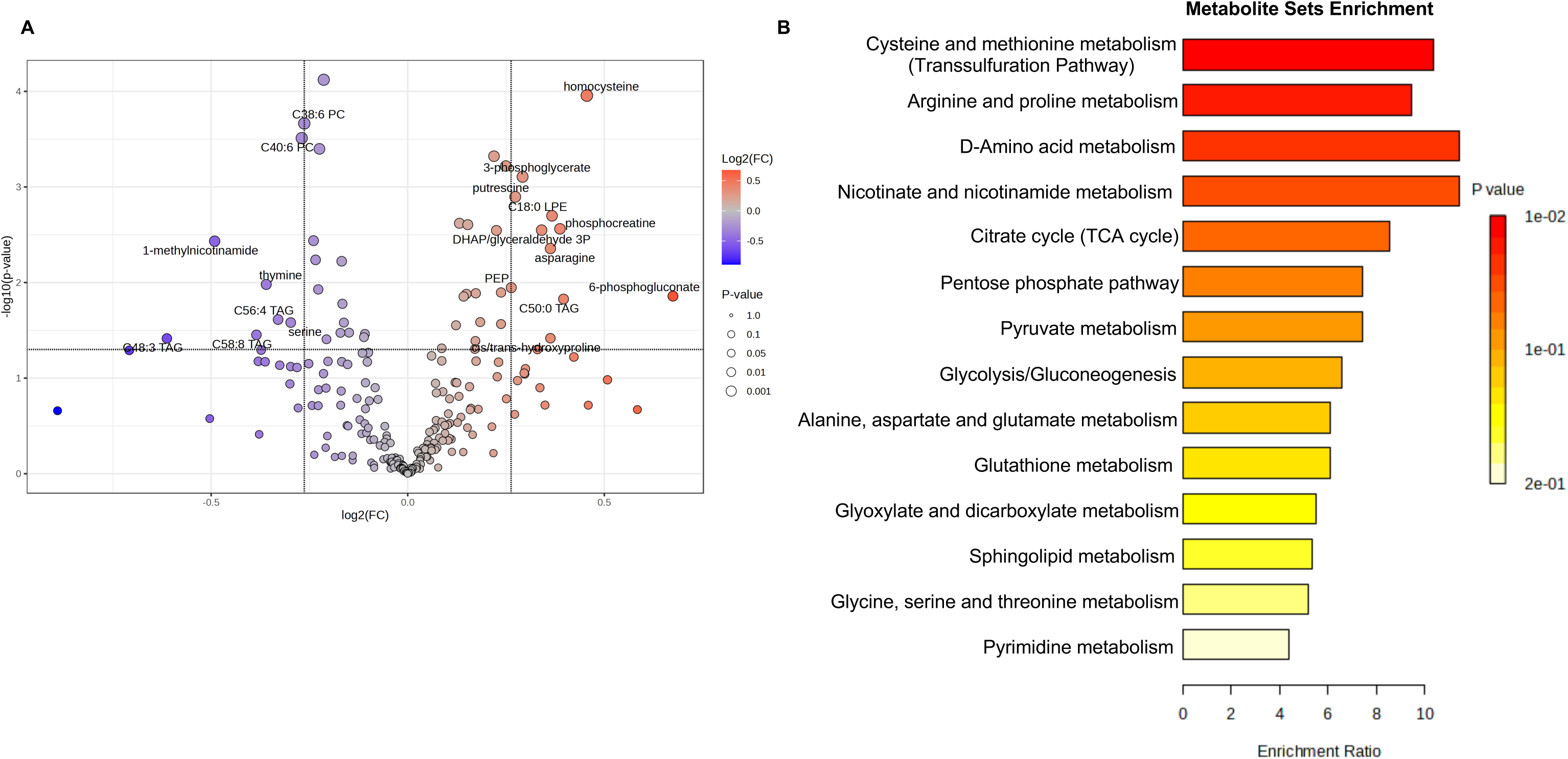
Profiling cancer metastasis-associated metabolites and metabolic pathways. **(A)** Volcano plot of metabolome was performed to compare the altered metabolites between metastatic cells with non-metastatic cells. Cutoff *p* value < 0.05; cutoff fold change (metastatic/non-metastatic) >1.2. **(B)** KEGG enrichment analysis of significantly altered metabolites in metastatic cells compared to non-metastatic cells (lipids are excluded). Unpaired *t*-test was performed.

To assess the functional relevance of these metabolic changes, we performed pathway enrichment analysis. As shown in **Figure 1B**, core metabolic pathways, such as glycolysis, glutathione metabolism, and the pentose phosphate pathway were highly enriched in metastatic cancer cells, consistent with previous reports [25–28]. Interestingly, additional pathways, including elevated cysteine and methionine metabolism (the transsulfuration pathway), arginine and proline metabolism, D-amino acid metabolism, nicotinate and nicotinamide metabolism, and the TCA cycle were also significantly associated with metastasis (**Figure 1B**). Collectively, these bioinformatic analyses provide an overview of metabolic alterations linked to cancer metastasis.

### Metastatic cancer cells display an organ-specific metabolic pattern

Cancer cells often metastasize to specific distant organs, including the liver, lung, bone, kidney, and brain [29]. To assess the role of metabolic alterations in driving organ colonization, we examined altered metabolic pathways in organ-specific metastasis (**Table S2**). Interestingly, metabolic differences were observed across different metastatic sites (**Figures 2** and **S1**). Specifically, for liver-specific metastases, elevated levels of sulfur-containing amino acids (e.g., cystathionine, homocysteine), nucleotides (e.g., CMP, GMP, AMP, UMP), and unique lipid species (e.g., C18:0 CE, C22:6 CE), and various TAGs were detected. In contrast, metabolites such as sorbitol, choline, and serine were significantly reduced (**Figure 2A**). The transsulfuration pathway (cysteine and methionine metabolism), glycine/serine/threonine metabolism, and purine metabolism were highly enriched for liver-specific metastasis (**Figure 2A**). For lung metastases, increases were observed in UMP, AMP, taurocholate, homocysteine, and lipid species such as C34:1 DAG and C18:0 LPE, while choline, phosphatidylcholines, and TAGs were downregulated. Pathway analysis revealed enrichment of glycine/serine/threonine metabolism, purine metabolism, and the transsulfuration pathway (cysteine and methionine metabolism) in lung-specific metastasis (**Figure 2B**). For brain metastasis, high levels of nucleotides, homocysteine, 6-phosphogluconate, and several lipid species were detected, while choline, phosphatidylcholines, and TAGs were found decreased (**Figure 2C**). Enriched metabolic pathways include pyrimidine and purine metabolism, the pentose phosphate pathway, and glycolysis (**Figure 2C**). For bone metastasis, elevated metabolites included homocysteine, taurocholate, 3-phosphoglycerate, glyceraldehyde-3P, and C18:0 LPE, whereas multiple lipid species, particularly TAGs and phosphatidylcholines, were reduced. Bone metastasis was characterized by increased fructose and mannose metabolism as well as taurine metabolism (**Figure 2D**). For kidney metastasis, increased metabolites included homocysteine, *N*-carbamoyl-beta-alanine, 6-phosphogluconate, and lipid species such as C14:0 CE and C22:6 CE. Conversely, thymine, uracil, and several TAGs (C56:4 TAG, C58:4 TAG, C54:4 TAG) were significantly decreased (**Figure 2E**). Pyrimidine metabolism, pantothenate and CoA biosynthesis, and β-alanine metabolism were significantly enriched in kidney metastasis (**Figure 2E**). Taken together, these findings suggest that metastatic cancer cells adopt distinct and organ-specific metabolic adaptations that likely facilitate their colonization within diverse microenvironments.

**Figure 2.**
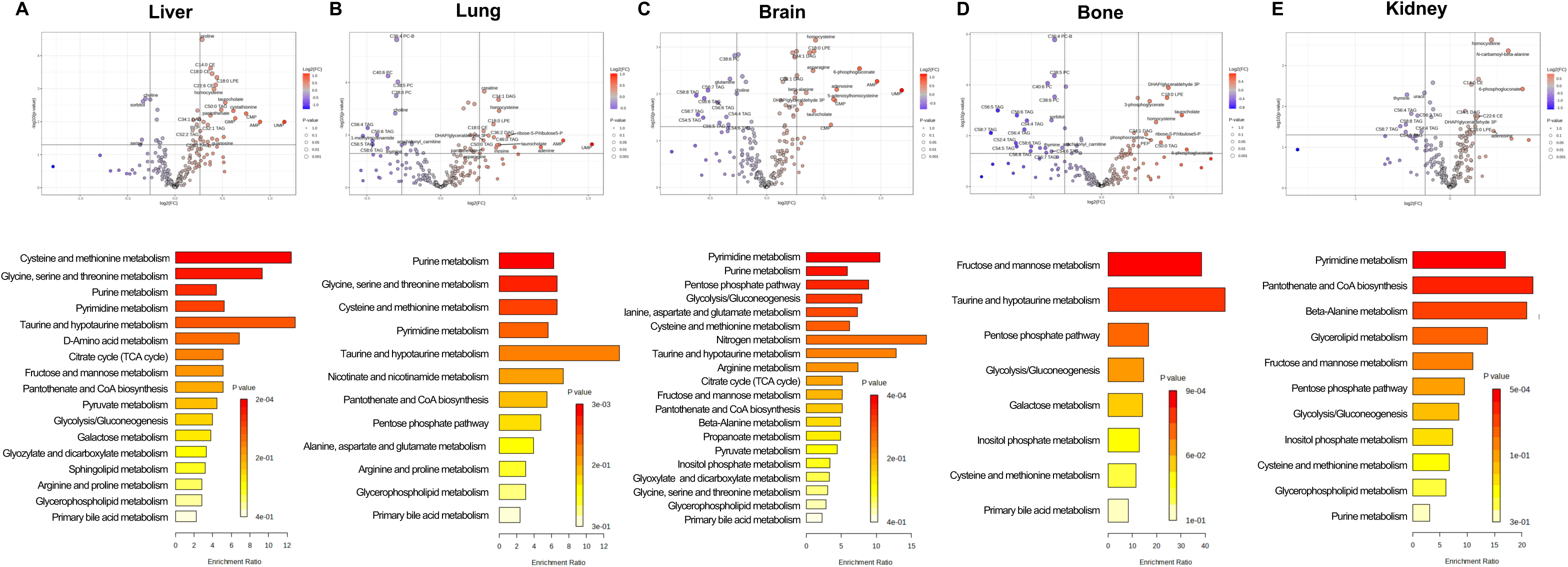
Cancer cells exert distinct metabolic landscapes across different metastatic sites. Volcano plot of metabolome combined with KEGG enrichment analysis was performed to compare altered metabolites and metabolic pathways between metastatic cell and non-metastatic cells metastasis to different organs, including (**A**) Liver, (**B**) Lung, (**C**) Brain, (**D**) Bone, (**E**) Kidney. Cutoff *p* value < 0.05; cutoff fold change (metastatic/non-metastatic) >1.2. Unpaired *t*-test was performed.

### Analysis of universal metabolites and metabolic pathways across different organ-specific metastases

Although different organ-specific metastases exhibit distinct metabolic features, our UpSet plot analysis [30] also revealed a set of shared metabolites (**Figure 3A** and **Table S3**). Among them, four metabolites (homocysteine, glyceraldehyde-3-P, lysophospholipid C18:0, and diacylglycerol C34:1) were consistently detected across all metastatic sites (**Figure 3A**). To uncover metabolic pathways most frequently shared among metastases, we conducted pathway enrichment analysis using metabolites present in at least three organ-specific metastatic sites. Interestingly, the transsulfuration pathway emerged as the most significantly enriched pathway (**Figure 3B**), consistent with our previous findings (**Figure 1B**). Additionally, pyrimidine metabolism, taurine metabolism, and the citrate cycle were also uncovered as common metabolic processes across different organ-specific metastases. Together, these results suggest the transsulfuration pathway as a central metabolic program that may drive metastatic progression across multiple organ sites.

**Figure 3.**
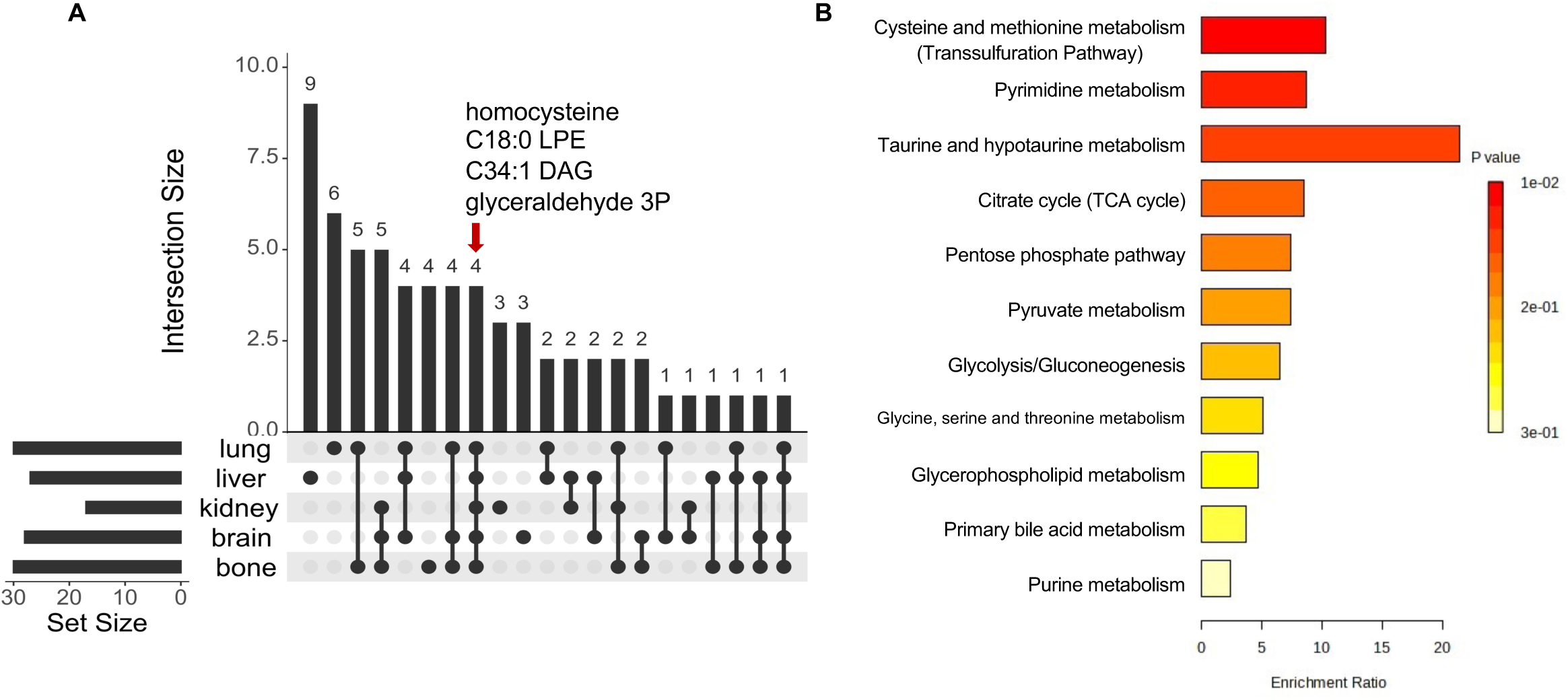
Analysis of shared metabolites and metabolic pathways across different organ-specific metastases. **(A)** UpSet plot analysis revealed the metabolites shared across indicated metastatic sites. Red arrow indicates the metabolites that are shared by all five organs. Cutoff *p* value < 0.05; cutoff fold change (metastatic/non-metastatic) >1.2. **(B)** Altered metabolites associated with at least 3 different organs were subjected to KEGG enrichment analysis. Unpaired *t*-test was performed.

### Cancer tissue of origin contributes to shaping the metabolic landscapes across different metastasis

Tissue of origin is known to influence organ-specific metastases [31], suggesting its potential role in shaping metastasis-associated metabolic reprogramming. To test this, we analyzed metabolic profiles of five highly metastatic cancers, including breast, skin, ovarian, colon, and pancreatic tumors (**Table S2**). Indeed, distinct tissue-dependent metabolic alterations were observed across these cancer types (**Figure 4**).

**Figure 4.**
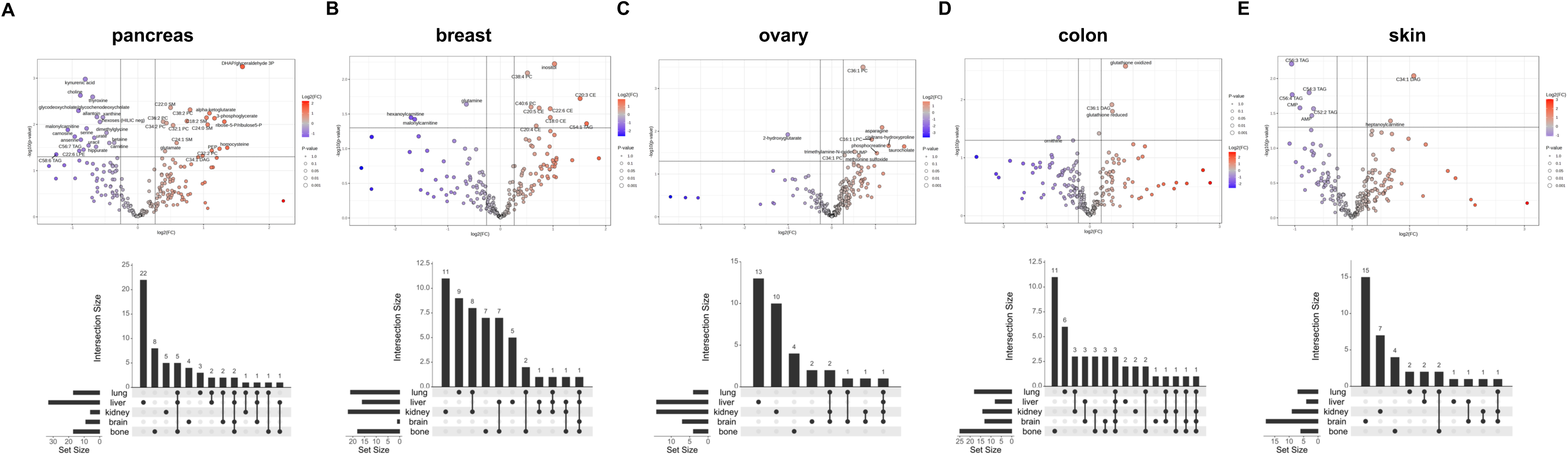
Cancer tissue of origin contributes to shaping the metabolic landscapes across different metastases. **(A)** Volcano plot of metabolome was performed to compare altered metabolites between metastatic cell with non-metastatic cells with different tissue of origin. Breast cancer, n=15 of non-metastatic group, n=7 in metastatic group; skin cancer, n=7 of non-metastatic group, n=32 in metastatic group; ovarian cancer, n=17 of non-metastatic group, n=15 in metastatic group; colon cancer, n=15 of non-metastatic group, n=12 in metastatic group; pancreatic cancer, n=8 of non-metastatic group, n=25 in metastatic group. **(B)** UpSet plot analysis revealed the metabolites shared across different metastatic sites from different cancer tissue of origin. Cutoff *p* value < 0.05; cutoff fold change (metastatic/non-metastatic) >1.2. Unpaired *t*-test was performed.

For pancreatic cancer, metabolites such as glyceraldehyde-3P, ribose-5-P, phosphoenolpyruvate, and homocysteine were significantly increased in metastatic cells, whereas choline, kynurenic acid, serine, betaine, and multiple TAG species were reduced. UpSet plot analysis showed the largest set of unique metabolites in liver metastases, suggesting a strong tissue-specific metabolic signature for pancreatic cancer. Across metastatic sites, 6-phosphogluconate and C32:1 PC were shared by four organs (**Figure 4A** and **Table S4**). For breast cancer, most upregulated metabolites in metastasis were lipid species, consistent with previous reports highlighting the role of lipids in breast cancer metastasis [32, 33]. In contrast, glutamine, hexanoylcarnitine, and malonylcarnitine were significantly downregulated, suggesting their potential tumor suppressive functions. UpSet analysis identified 11 and 9 unique metabolites associated with kidney and lung metastases, respectively (**Figure 4B** and **Table S4**). Among the identified metabolites, lactose was shared across metastatic sites (**Figure 4B** and **Table S4**), suggesting its potential as a biomarker for breast cancer metastasis. For ovarian cancer, C36:1 PC, asparagine, taurocholate, and phosphocreatine were significantly increased in metastatic cells, while 2-hydroxyglutarate was downregulated. Most altered metabolites were associated with liver metastases, followed by kidney metastases. Arginine was the only metabolite shared across four different metastatic sites (**Figure 4C** and **Table S4**). For colorectal cancer, elevated glutathione (oxidized or reduced) and C36:1 DAG, along with reduced ornithine, were associated with overall metastasis. We identified 11 metabolites linked to bone metastases and 6 to lung metastases. Among them, lactate, malondialdehyde, and ribose-5-P were shared across all metastasis sites (**Figure 4D** and **Table S4**). For skin cancer, hepanoylcarnitine and C34:1 DAG were upregulated in metastatic cells, while CMP, AMP, and several TAG species were downregulated. Organ-specific analysis revealed 15 metabolites associated with brain metastases and 7 with kidney metastases, with alpha-hydroxybutyrate shared across three metastatic sites (**Figure 4E** and **Table S4**).

Collectively, these data suggest that tissue of origin modulates metabolic programs that shape the metastatic landscape and influence organ-specific colonization.

### The transsulfuration pathway is activated in metastatic pancreatic cancer cells

Pancreatic ductal adenocarcinoma (PDAC) is a highly metastatic disease with a 5-year survival rate of less than 7% [34]. More than half of PDAC cases present with liver metastasis [35]. Notably, our bioinformatic analyses revealed the transsulfuration pathway as one of the most enriched pathways in liver metastasis (**Figure 2A**). Its key metabolite, homocysteine, was significantly associated with pancreatic cancer metastasis (**Figure 4A**). These findings together suggest an oncogenic role for this metabolic pathway in PDAC progression.

The transsulfuration pathway, also known as the cysteine and methionine pathway, converts homocysteine to cysteine via the enzymes cystathionine β-synthase (CBS) and cystathionine γ-lyase (CGL/CTH) (**Figure 5A**). The Cancer Genome Atlas (TCGA) analysis showed that high CGL expression strongly correlated with poor overall survival of PDAC patients (**Figure 5B**), underscoring the pathological relevance of dysregulated transsulfuration pathway activity. Consistently, both CBS and CGL were highly expressed in metastatic PDAC cell lines PATU8988 and SUIT2 compared to non-metastatic cell lines HUPT3 and CAPAN2 (**Figure 5C**). Moreover, levels of methionine, cystathionine and cysteine were dramatically increased in metastatic cells as compared to non-metastatic cells (**Figure 5D**). As a control, methionine deprivation abolished the increase of cystathionine and cysteine in metastatic PDAC cells (**Figure 5D**), confirming the involvement of the transsulfuration pathway in this process. Moreover, a hierarchical clustering heatmap revealed distinct metabolic signatures between non-metastatic (HUPT3 and CAPAN-2) and metastatic (SUIT2 and PATU8988) pancreatic cancer cell. Under control conditions, metastatic cells displayed higher levels of metabolites involved in the transsulfuration pathway and tricarboxylic acid (TCA) cycles, such as cystathionine, succinate, fumarate, malate, methionine, and cysteine. Interestingly, methionine deprivation leads to suppression of these metabolites to levels similar to non-metastatic cells (**Figure S2A**). To determine the overall redox balance with increased sulfur metabolism, we analyzed GSH/GSSG ratio and found no significant difference while homocysteine level is increased (**Figure S2B**). Taken together, these data show that the transsulfuration pathway is highly active in metastatic pancreatic cancer cells and may contribute to their metastasis.

**Figure 5.**
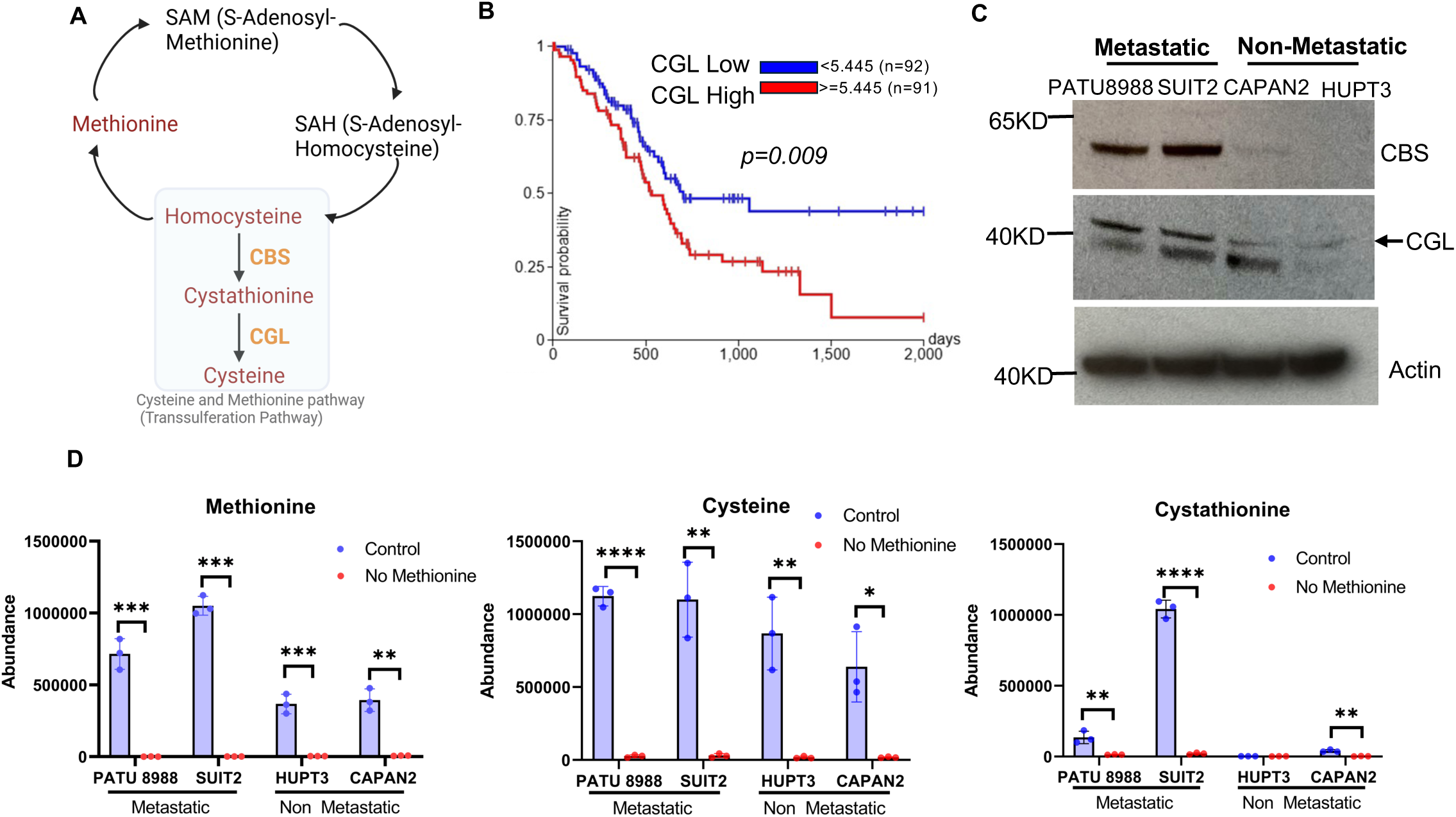
The transsulfuration pathway activity is highly increased in metastatic pancreatic cancer cells. **(A)** Schematic illustration of the transsulfuration pathway. Two key enzymes CBS and CGL are shown. **(B)** Kaplan-Meier curves of overall survival of patients with pancreatic cancer is stratified by *CGL* expression level. **(C)** Western blot was performed using the indicated antibodies. **(D)** GC-MS analyses were performed for the indicated metabolites in different pancreatic cancer cells. Cells were treated with methionine deprivation for 0 and 24 hours (n=3). Unpaired *t*-test was performed. \*\**p* < 0.01, \*\*\**p* < 0.001, \*\*\*\**p* < 0.0001.

### The transsulfuration pathway is required for metastatic pancreatic cancer cell migration and invasion

Given the association between the transsulfuration pathway and PDAC metastasis (**Figures 5C-D**), we next examined its role in this malignant process. Wound healing assays showed that methionine deprivation significantly suppressed the migration of three metastatic PDAC cell lines, PATU8988 (**Figure 6A**), SUIT2 (**Figure 6A**), and PANC-1 (**Figure S3**). Similarly, cell invasion assays demonstrated that methionine deprivation dramatically impaired SUIT2 cell invasion (**Figure 6B**) without affecting the overall cell viability (**Figure 6C**).

**Figure 6.**
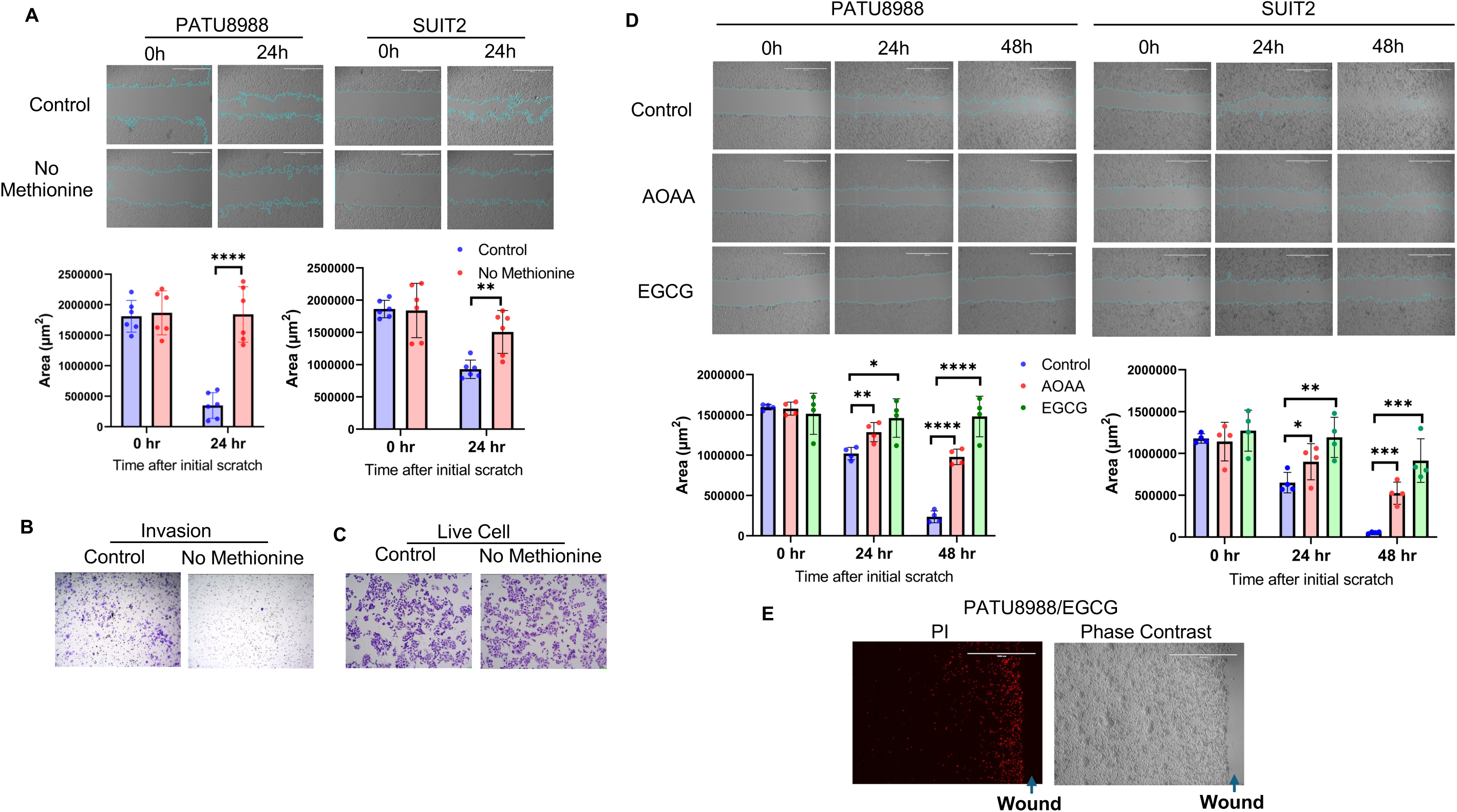
The transsulfuration pathway is required for metastatic pancreatic cancer cell migration and invasion. **(A)** Wound healing assays were performed using PATU8988 and SUIT2 cells under control and methionine deprivation conditions. Wound area was quantified using Image J software (n=6). Representative images were shown. Unpaired *t*-test was performed. \*\**p* < 0.01, \*\*\*\**p* < 0.0001. Scale bar, 1000 μm. **(B)** Transwell invasion assays were performed using SUIT2 cells under control and methionine deprivation conditions. **(C)** Methionine deprivation did not affect SUIT2 cell viability. Crystal violet staining was performed for the cells in transwell assays (**B**). **(D)** Pharmacological inhibition of CBS suppressed PATU8988 and SUIT2 cell migration. Cells were treated with AOAA (10 mM) or EGCG (0.1 mM) for 0, 24 and 48 hours. Wound area was quantified using Image J software (n=4). Representative images were shown. Unpaired *t*-test was performed. \**p* < 0.05, \*\**p* < 0.01, \*\*\**p* < 0.001, \*\*\*\**p* < 0.0001. Scale bar, 1000 μm. **(E)** Representative pictures of PATU8988 cell, treated with EGCG (0.1mM, 48 hours) and stained with propidium iodide (PI) (left panel). The same area was shown with the phase contrast (right panel).

To further validate these findings, we pharmacologically inhibited the transsulfuration pathway using two established CBS inhibitors: aminooxyacetic acid (AOAA), a commonly used CBS inhibitor [36], and epigallocatechin gallate (EGCG), the major bioactive component of green tea and a potent CBS inhibitor [37]. Consistently, both AOAA and EGCG significantly reduced metastatic PDAC cell migration (**Figure 6D**). Although EGCG treatment did not affect overall cell viability (**Figure 6E**), propidium iodide (PI) staining revealed specific cell death of migratory cells in wound healing assays (**Figure 6E**), suggesting a potential mechanism by which CBS inhibition selectively targets migratory populations.

Collectively, these findings demonstrate an essential role of the transsulfuration pathway for PDAC cell migration and invasion and suggest it as a potential therapeutic target for metastatic PDAC.

## Discussion

Metastasis is known as the leading cause of cancer-related mortality, yet the mechanisms underlying this process, particularly the contribution of metabolic reprogramming, are still largely unknown. While prior studies have independently profiled cancer cell metabolism and metastatic potential, our integrative analysis bridges these two datasets to systematically define the metabolites and metabolic pathways associated with metastatic cancer cells, thereby providing a global view of metastasis-related metabolic reprogramming.

Our bioinformatic analyses uncover distinct metabolic alterations across different cancers and their organ-specific metastases, supporting the “seed and soil” hypothesis. Importantly, we demonstrate that the tissue of origin exerts critical influence on the metabolic landscape of metastases, consistent with recent evidence that primary tumor identity shapes both primary and metastatic tumor metabolisms to impact its organ-specific metastasis [38]. Notably, metastatic tumors also retain metabolic signatures inherited from their tissue of origin, while adapting to the metabolic demands of their new microenvironments. This dual influence contributes to the heterogeneity of metabolic phenotypes observed across different metastatic niches. Our findings thus underscore the critical role of metabolism in driving metastasis. Dissecting the metabolic dependencies of metastatic cells will not only enrich our mechanistic understanding of cancer metastasis, but may also uncover therapeutic vulnerabilities that can be exploited in a tissue- and site-specific manner.

Interestingly, our study identified the transsulfuration pathway as one of the most enriched metabolic alterations in metastatic cancer cells. This pathway transfers sulfur from homocysteine to cysteine via cystathionine and serves as the only metabolic route for cysteine biosynthesis [39]. The process is mediated by two key enzymes, CBS and CGL/CTH. Notably, CBS is highly expressed in multiple cancers [40] and has been shown to promote cancer metastasis [41, 42], and CGL/CTH is known to promote prostate and breast cancer metastases [43] [44]. A recent study using multiomic screening of invasive glioblastoma cells reveals that CGL/CTH is essential for glioblastoma invasion [45]. Consistent with these findings, our data show that both CBS and CGL are elevated in metastatic pancreatic cancer cells (**Figure 5C**) and that increased CBS expression correlates with poor patient survival in pancreatic cancer (**Figure 5B**). Furthermore, inhibition of the transsulfuration pathway through either methionine deprivation or treatment with CBS inhibitors suppressed migration and invasion of metastatic pancreatic cancer cells (**Figures 6A-D**). Although these results highlight an oncogenic role for the transsulfuration pathway in pancreatic cancer metastasis, the underlying mechanisms remain to be fully addressed.

One potential mechanism involves cysteine, the downstream metabolite of the transsulfuration pathway. Cysteine serves as a critical substrate for glutathione synthesis, one of the most important intracellular antioxidants. This dependency suggests that the transsulfuration pathway is essential for metastatic cancer cells, where glutathione plays a critical role in reducing oxidative stress induced by reactive oxygen species (ROS) [46]. Although low ROS levels can promote cell proliferation, migration, and invasion, excessive ROS accumulation resulting from dysregulated metabolic activity and redox homeostasis can damage DNA, lipids, and proteins, leading to cell death [47, 48]. To survive, cancer cells thus have to fine-tune ROS levels, and our data suggest that the transsulfuration pathway is likely to contribute to this regulation by supplying cysteine and supporting glutathione biosynthesis. In line with this notion, inhibition of the transsulfuration pathway induced apoptosis in migratory pancreatic cancer cells (**Figure 6E**). Future mechanistic studies are needed to assess the contribution of glutathione-mediated ROS clearance to this process. Moreover, accumulating evidence supports that the metabolites from the transsulfuration pathway, such as hydrogen sulfide (H_2_S), play a role in epigenetic regulation via modifying histone proteins sulfhydration [49]. H_2_S has been shown to regulate the activity of DNA methyltransferases (DNMTs), the enzymes responsible for DNA methylation [50]. It has been well acknowledged that epigenetic change is involved in cancer metastasis [51]. Therefore, it is possible that an altered transsulfuration pathway may affect metastasis via epigenetic regulation. Moreover, H₂S functions as a signaling mediator affecting stromal and immune cells in the tumor microenvironment, suggesting another mechanism by which transsulfuration pathway affects metastasis [52]. Future work is needed to study epigenetic and microenvironmental remodeling to elucidate the underlying mechanism. For example, *in vivo* metabolic tracing using isotope-labeled precursors such as ^13^C-methionine could be employed to quantify flux through the transsulfuration pathway in metastatic models. In addition, inhibiting key transsulfuration enzymes (CBS, CTH/CGL) through genetic manipulation in cancer cells would help further confirm the role of pathway activity in modulating epigenetics, tumor microenvironment, and metastatic phenotypes.

Our study also highlights the translational significance of targeting the transsulfuration pathway for treating cancer metastasis. We demonstrate that methionine deprivation is effective in reducing intracellular cystathionine in pancreatic cancer cells (**Figure 5D**). While methionine metabolism has been previously implicated in driving cancer cell survival and epigenetic regulation [53], its association with metastasis has not been fully investigated. Notably, dietary methionine restriction is sufficient to reduce circulating methionine in most human and animal studies within just days of consuming such diets [54]. Together with our findings, this evidence suggests that the methionine restriction diet could be used to treat or prevent cancer metastasis. Moreover, as a potent CBS inhibitor, EGCG is an over-the-counter dietary supplement extracted from green tea that may help with weight management and anti-inflammation [55, 56]. Although EGCG has been shown to have beneficial effects on pancreatic cancer chemotherapeutic responses [57] or inhibit pancreatic cancer cell growth [58], its inhibitory effect on pancreatic cancer metastasis has not been explored. According to our findings (**Figure 6D-E**), EGCG holds the potential to treat or prevent pancreatic cancer metastasis, supporting the anti-cancer effect of green tea. Despite these interesting findings, it will be highly important to further investigate the target role of the transsulfuration pathway as well as the therapeutic potentials of the methionine restriction diet and EGCG against pancreatic cancer metastasis using *in vivo* animal models.

Several limitations are noticed for our current study. First, metastasis potential was simplified to a binary variable (0 or 1), which may not fully represent the accuracy of metastatic behavior. Moreover, although volcano plot analysis has revealed that lipid metabolism plays a crucial role in metastasis, our current findings have not been able to uncover the pathways for the altered lipid metabolism. Next, our current experimental studies were mostly performed *in vitro*, which cannot fully address the critical influence of the tumor microenvironment on metastatic processes. Additional *in vivo* studies are needed to further strengthen the translational significance of our work. Finally, the molecular mechanisms by which altered transsulfuration pathway regulates cancer metastasis remain to be addressed in the future.

## Conclusion

In this study, we systematically profiled metabolic alterations between metastatic and non-metastatic cancer cells by integrating published MetMap datasets and CCLE metabolomics data. Our results not only indicate that distinct metabolic pathways are associated with organ-specific metastasis, but also highlight the crucial role of the tissue of origin in shaping the metabolic landscape of metastatic tumors. Moreover, we uncovered several metabolic pathways highly associated with metastasis. Among them, the transsulfuration pathway was found essential for metastatic pancreatic cancer cell migration and invasion, underscoring its target role for treating metastatic cancers.

## Acknowledgements

We thank Dr. Christopher Halbrook (University of California, Irvine) for providing the pancreatic cancer cell lines. This work was supported by the Setup funds provided to W.W. by the School of Biological Sciences at University of California, Irvin. Research reported in this publication was also supported in part by NIH-NCI grant (P30CA062203) and the UCI Chao Family Comprehensive Cancer Center using Anti-Cancer Challenge funds.

## Author contributions

J.K.Y. conceived the study, performed the bioinformatic analyses and experiments, and wrote the manuscript. Y.Y. conducted the GC-MS analysis. W.W. edited the manuscript.

## Key points

- Bioinformatic analysis defines the metabolic landscape associated with cancer metastasis.
- Organ-specific metastasis involves distinct metabolites and metabolic pathways.
- Tissue of origin contributes to the metabolic features of metastatic tumors.
- The transsulfuration pathway is highly activated in metastatic pancreatic cancer cells and essential for cell migration and invasion.

---

**Figure S1.**
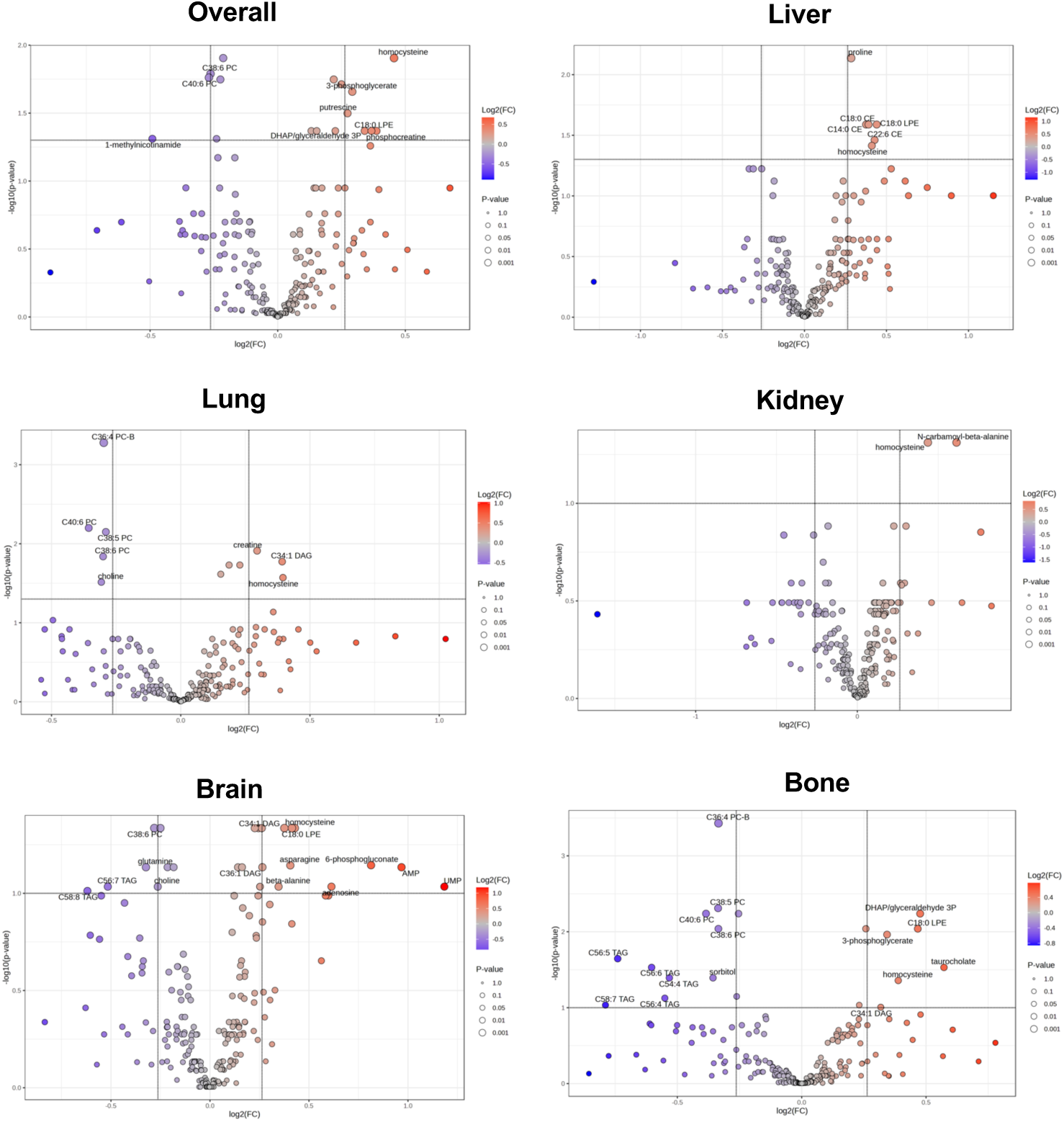
Volcano plot of the metabolome comparing altered metabolites between metastatic and non-metastatic cells. Significance was assessed using an FDR-adjusted p-value threshold < 0.05, with a fold-change cutoff (metastatic/non-metastatic) > 1.2.

**Figure S2.**
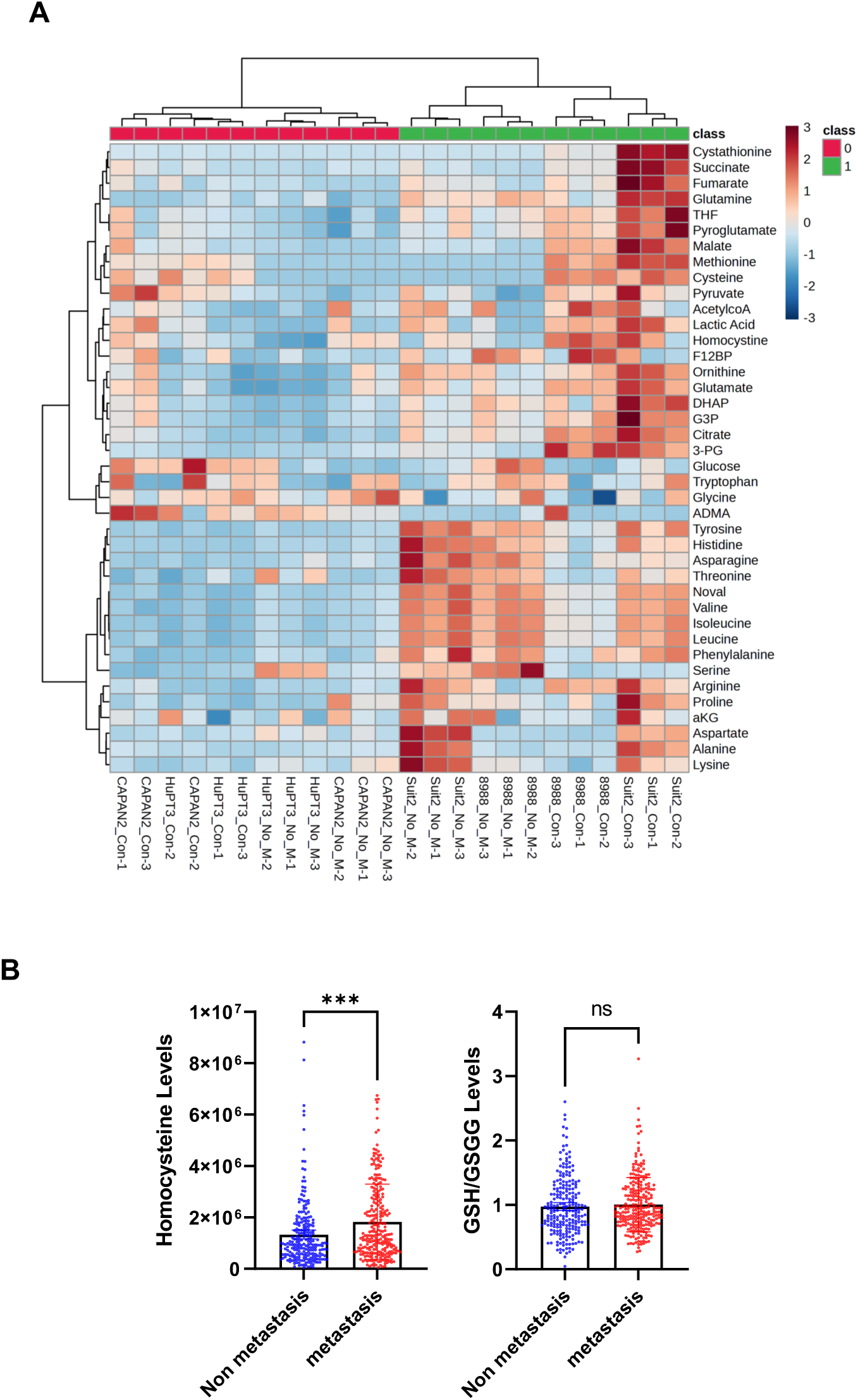
**A.** Heatmap of significantly changed metabolites in CAPAN2, HuPT3, Suit2 and Patu8988 from control media (Con) and methionine deprived media (No M) for 24 hours by GC-MS (n=3 per group). B. Comparison of the indicated metabolites between metastatic cells (overall metastatic potential) with non-metastatic cells (n=223 for non metastatic, n=256 for metastatic cells). *p < 0.05, ***p < 0.001, ns, non significant, by unpaired t-test.

**Figure S3.**
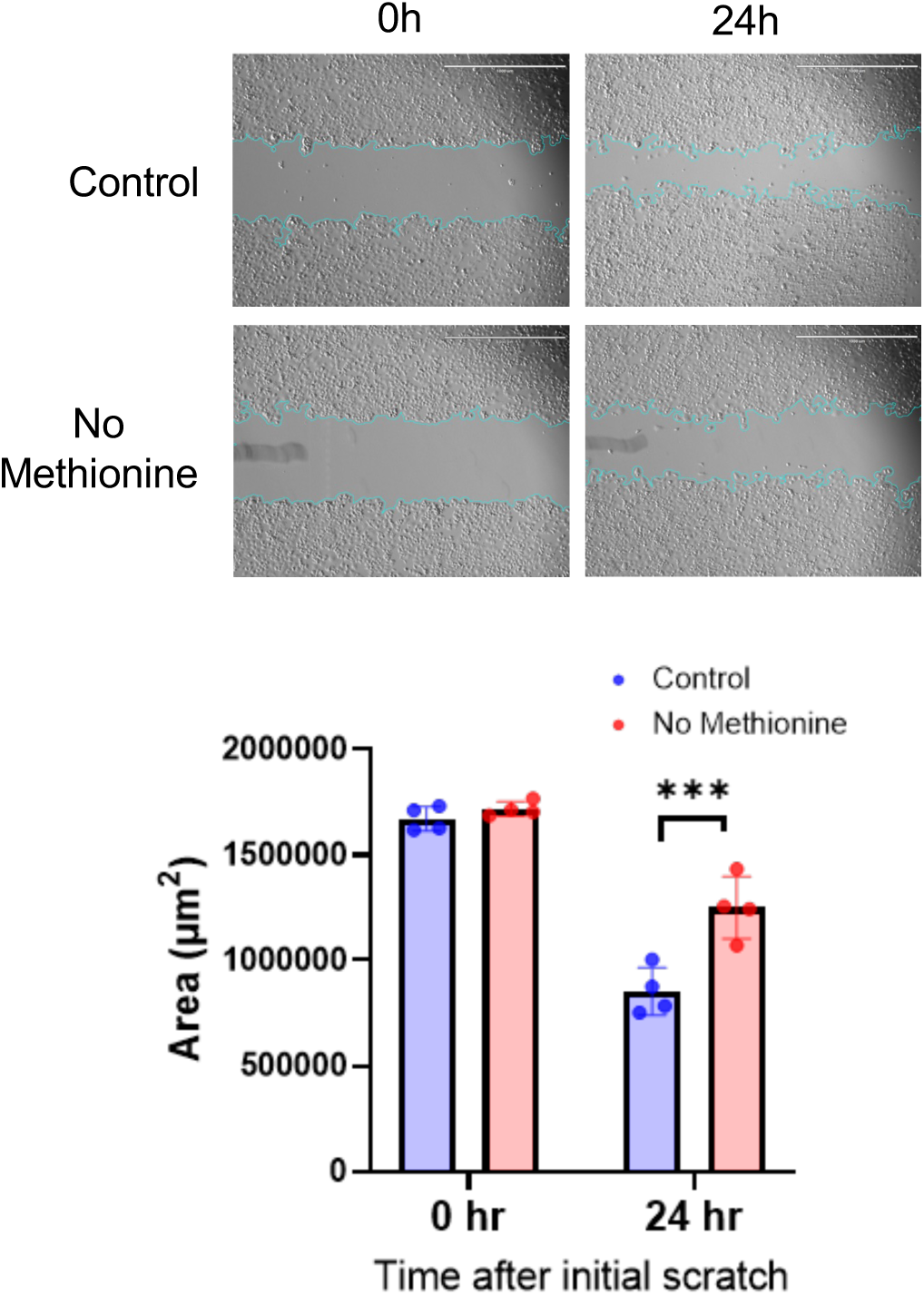
Wound healing assays were performed using Panc-1 cell under control and methionine deprivation conditions. Wound area was quantified using Image J software (n=6). Representative images were shown. Unpaired t-test was performed. ***p < 0.001. Scale bar, 1000 μm.

## References

1. Ferlay J, Colombet M, Soerjomataram I et al. Cancer statistics for the year 2020: An overview, Int J Cancer 2021.

2. Chaffer CL, Weinberg RA. A perspective on cancer cell metastasis, Science 2011;331:1559–1564.

3. Dai X, Xi M, Li J. Cancer metastasis: molecular mechanisms and therapeutic interventions, Mol Biomed 2025;6:20.

4. Gupta GP, Massague J. Cancer metastasis: building a framework, Cell 2006;127:679–695.

5. Talmadge JE, Fidler IJ. AACR centennial series: the biology of cancer metastasis: historical perspective, Cancer Res 2010;70:5649–5669.

6. Li Y, Liu F, Cai Q et al. Invasion and metastasis in cancer: molecular insights and therapeutic targets, Signal Transduct Target Ther 2025;10:57.

7. van Zijl F, Krupitza G, Mikulits W. Initial steps of metastasis: cell invasion and endothelial transmigration, Mutat Res 2011;728:23–34.

8. Fidler IJ. The pathogenesis of cancer metastasis: the ‘seed and soil’ hypothesis revisited, Nat Rev Cancer 2003;3:453–458.

9. Akhtar M, Haider A, Rashid S et al. Paget’s “Seed and Soil” Theory of Cancer Metastasis: An Idea Whose Time has Come, Adv Anat Pathol 2019;26:69–74.

10. Zhao K, Vos J, Lam S et al. Longitudinal and multisite sampling reveals mutational and copy number evolution in tumors during metastatic dissemination, Nat Genet 2025;57:1504–1511.

11. Murcott B, Honig F, Halliwell DO et al. Colorectal cancer progression to metastasis is associated with dynamic genome-wide biphasic 5-hydroxymethylcytosine accumulation, BMC Biol 2025;23:100.

12. Hu D, Zhao T, Xu C et al. Epigenetic Modifiers in Cancer Metastasis, Biomolecules 2024;14.

13. Punnasseril JMJ, Auwal A, Gopalan V et al. Metabolic Reprogramming of Cancer Cells and Therapeutics Targeting Cancer Metabolism, Cancer Med 2025;14:e71244.

14. DeBerardinis RJ, Chandel NS. We need to talk about the Warburg effect, Nat Metab 2020;2:127–129.

15. Payne KK. Cellular stress responses and metabolic reprogramming in cancer progression and dormancy, Semin Cancer Biol 2022;78:45–48.

16. Gounis M, Campos AV, Shokry E et al. Metabolic adaptations of micrometastases alter EV production to generate invasive microenvironments, J Cell Biol 2025;224.

17. Elia I, Doglioni G, Fendt SM. Metabolic Hallmarks of Metastasis Formation, Trends Cell Biol 2018;28:673–684.

18. Teoh ST, Lunt SY. Metabolism in cancer metastasis: bioenergetics, biosynthesis, and beyond, Wiley Interdiscip Rev Syst Biol Med 2018;10.

19. Faubert B, Solmonson A, DeBerardinis RJ. Metabolic reprogramming and cancer progression, Science 2020;368.

20. Jin X, Demere Z, Nair K et al. A metastasis map of human cancer cell lines, Nature 2020;588:331–336.

21. Li H, Ning S, Ghandi M et al. The landscape of cancer cell line metabolism, Nat Med 2019;25:850–860.

22. Chong J, Xia J. MetaboAnalystR: an R package for flexible and reproducible analysis of metabolomics data, Bioinformatics 2018;34:4313–4314.

23. Conway JR, Lex A, Gehlenborg N. UpSetR: an R package for the visualization of intersecting sets and their properties, Bioinformatics 2017;33:2938–2940.

24. Suarez-Arnedo A, Torres Figueroa F, Clavijo C et al. An image J plugin for the high throughput image analysis of in vitro scratch wound healing assays, PLoS One 2020;15:e0232565.

25. Wei Q, Qian Y, Yu J et al. Metabolic rewiring in the promotion of cancer metastasis: mechanisms and therapeutic implications, Oncogene 2020;39:6139–6156.

26. Piskounova E, Agathocleous M, Murphy MM et al. Oxidative stress inhibits distant metastasis by human melanoma cells, Nature 2015;527:186–191.

27. Biondini M, Lehuede C, Tabaries S et al. Metastatic breast cancer cells are metabolically reprogrammed to maintain redox homeostasis during metastasis, Redox Biol 2024;75:103276.

28. Patra KC, Hay N. The pentose phosphate pathway and cancer, Trends Biochem Sci 2014;39:347–354.

29. Obenauf AC, Massague J. Surviving at a Distance: Organ-Specific Metastasis, Trends Cancer 2015;1:76–91.

30. Lex A, Gehlenborg N, Strobelt H et al. UpSet: Visualization of Intersecting Sets, IEEE Trans Vis Comput Graph 2014;20:1983–1992.

31. Carrolo M, Miranda JAI, Vilhais G et al. Metastatic organotropism: a brief overview, Front Oncol 2024;14:1358786.

32. Martin-Perez M, Urdiroz-Urricelqui U, Bigas C et al. The role of lipids in cancer progression and metastasis, Cell Metab 2022;34:1675–1699.

33. Fu W, Sun A, Dai H. Lipid metabolism involved in progression and drug resistance of breast cancer, Genes Dis 2025;12:101376.

34. Allemani C, Matsuda T, Di Carlo V et al. Global surveillance of trends in cancer survival 2000-14 (CONCORD-3): analysis of individual records for 37 513 025 patients diagnosed with one of 18 cancers from 322 population-based registries in 71 countries, Lancet 2018;391:1023–1075.

35. Liu Z, Gou A, Wu X. Liver metastasis of pancreatic cancer: the new choice at the crossroads, Hepatobiliary Surg Nutr 2023;12:88–91.

36. Petrosino M, Zuhra K, Kopec J et al. H(2)S biogenesis by cystathionine beta-synthase: mechanism of inhibition by aminooxyacetic acid and unexpected role of serine, Cell Mol Life Sci 2022;79:438.

37. Zuhra K, Petrosino M, Gupta B et al. Epigallocatechin gallate is a potent inhibitor of cystathionine beta-synthase: Structure-activity relationship and mechanism of action, Nitric Oxide 2022;128:12–24.

38. Sivanand S, Gultekin Y, Winter PS et al. Cancer tissue of origin constrains the growth and metabolism of metastases, Nat Metab 2024;6:1668–1681.

39. Sbodio JI, Snyder SH, Paul BD. Regulators of the transsulfuration pathway, Br J Pharmacol 2019;176:583–593.

40. Ascencao K, Szabo C. Emerging roles of cystathionine beta-synthase in various forms of cancer, Redox Biol 2022;53:102331.

41. Czikora A, Erdelyi K, Ditroi T et al. Cystathionine beta-synthase overexpression drives metastatic dissemination in pancreatic ductal adenocarcinoma via inducing epithelial-to-mesenchymal transformation of cancer cells, Redox Biol 2022;57:102505.

42. Liu Y, Pan L, Li Y et al. Cystathionine-beta-synthase (CBS)/H2S system promotes lymph node metastasis of esophageal squamous cell carcinoma (ESCC) by activating SIRT1, Carcinogenesis 2022;43:382–392.

43. Wang YH, Huang JT, Chen WL et al. Dysregulation of cystathionine gamma-lyase promotes prostate cancer progression and metastasis, EMBO Rep 2019;20:e45986.

44. Wang L, Shi H, Liu Y et al. Cystathionine-gamma-lyase promotes the metastasis of breast cancer via the VEGF signaling pathway, Int J Oncol 2019;55:473–487.

45. Garcia JH, Akins EA, Jain S et al. Multiomic screening of invasive GBM cells reveals targetable transsulfuration pathway alterations, J Clin Invest 2023;134.

46. Kwon DH, Cha HJ, Lee H et al. Protective Effect of Glutathione against Oxidative Stress-induced Cytotoxicity in RAW 264.7 Macrophages through Activating the Nuclear Factor Erythroid 2-Related Factor-2/Heme Oxygenase-1 Pathway, Antioxidants (Basel) 2019;8.

47. Chen D, Guo Z, Yao L et al. Targeting oxidative stress-mediated regulated cell death as a vulnerability in cancer, Redox Biol 2025;84:103686.

48. Nakamura H, Takada K. Reactive oxygen species in cancer: Current findings and future directions, Cancer Sci 2021;112:3945–3952.

49. Chen HJ, Qian L, Li K et al. Hydrogen sulfide-induced post-translational modification as a potential drug target, Genes Dis 2023;10:1870–1882.

50. Dogaru BG, Munteanu C. The Role of Hydrogen Sulfide (H(2)S) in Epigenetic Regulation of Neurodegenerative Diseases: A Systematic Review, Int J Mol Sci 2023;24.

51. Janin M, Davalos V, Esteller M. Cancer metastasis under the magnifying glass of epigenetics and epitranscriptomics, Cancer Metastasis Rev 2023;42:1071–1112.

52. Machado-Neto JA, Cerqueira ARA, Verissimo-Filho S et al. Hydrogen Sulfide Signaling in the Tumor Microenvironment: Implications in Cancer Progression and Therapy, Antioxid Redox Signal 2024;40:250–271.

53. Sanderson SM, Gao X, Dai Z et al. Methionine metabolism in health and cancer: a nexus of diet and precision medicine, Nat Rev Cancer 2019;19:625–637.

54. Gao X, Sanderson SM, Dai Z et al. Dietary methionine influences therapy in mouse cancer models and alters human metabolism, Nature 2019;572:397–401.

55. Yan R, Cao Y. The Safety and Efficacy of Dietary Epigallocatechin Gallate Supplementation for the Management of Obesity and Non-Alcoholic Fatty Liver Disease: Recent Updates, Biomedicines 2025;13.

56. Chen IJ, Liu CY, Chiu JP et al. Therapeutic effect of high-dose green tea extract on weight reduction: A randomized, double-blind, placebo-controlled clinical trial, Clin Nutr 2016;35:592–599.

57. Tang SN, Fu J, Shankar S et al. EGCG enhances the therapeutic potential of gemcitabine and CP690550 by inhibiting STAT3 signaling pathway in human pancreatic cancer, PLoS One 2012;7:e31067.

58. Bimonte S, Leongito M, Barbieri A et al. Inhibitory effect of (-)-epigallocatechin-3-gallate and bleomycin on human pancreatic cancer MiaPaca-2 cell growth, Infect Agent Cancer 2015;10:22.

